# A Mesoscale Finite Element Modelling Approach for Understanding Brain Morphology and Material Heterogeneity Effects in Chronic Traumatic Encephalopathy

**DOI:** 10.1101/2020.06.09.141259

**Authors:** A. Bakhtairydavijani, G. Khalid, M.A. Murphy, K.L. Johnson, L. E. Peterson, M. Jones, M.F. Horstemeyer, A.C. Dobbins, R. K. Prabhu

**Author notes:** Corresponding Author: Assistant Professor, Department of Agricultural and Biological Engineering, 130 Creelman St, Mississippi State University, MS 39762.

## Abstract

Chronic Traumatic Encephalopathy (CTE) affects a significant portion of athletes in contact sports but is difficult to quantify using clinical examinations and modelling approaches. We use an *in silico* approach to quantify CTE biomechanics using mesoscale Finite Element (FE) analysis that bridges with macroscale whole head FE analysis. The sulci geometry produces complex stress waves that interact with each another to create increased shear stresses at the sulci depth that are significantly larger than in analyses without sulci (from 0.5 kPa to 18.0 kPa). Also, Peak sulci stresses are located where CTE has been experimentally observed in the literature.

**Highlights:** 3 to 5 bullet points 85 characters max

- Sulci introduce stress localizations at their depth in the gray matter
- Sulci stress fields interact to produce stress concentration sites in white matter
- Differentiating brain tissue properties did not significantly affect peak stresses

## Introduction

Chronic Traumatic Encephalopathy (CTE) is a progressive neurodegenerative disease common to a range of contact sports (Bailes et al. 2013). Parker (Parker 1934) first documented CTE, also known as punch drunk syndrome or dementia pugilistica, in boxers. This neurodegeneration has since been identified in multiple sports, including boxing (Saing et al. 2012), wrestling (Cajigal 2007), soccer (Geddes et al. 1999), and American professional football (Omalu et al. 2005). More recently, a post-mortem study of National Football League (NFL) players found 110 out of 111 of the brains examined suffered from CTE (Mez et al. 2017). Considering the progressive nature of this disease, and the lack of sufficient protection through protective gear and guidelines, further study of this disease and its underlying causes can help identify preventive measures and thus benefit the quality of life of these individuals.

Significant evidence points to the relationship between multiple sub-concussions to mild Traumatic Brain Injury (mTBI) and chronic neurological injury. American football players are estimated to experience 100 to 1000 head impacts in a season (Bailes et al. 2013) where more than three concussions correlate with an increased probability of mild cognitive impairment and depression (Guskiewicz et al. 2007). At the same time, it results in a greater probability of future concussions with longer recovery times (McCrea et al. 2003). The CTE pathology includes deposits of tau protein in the form of NeuroFibrillary Tangles (NFTs) and astrocytic tangles, preferentially distributed in superficial layers (Layer II and upper third of layer III) in neocortical areas (Hof et al. 1991); amyloid-β plaques (Corsellis et al. 1973); and vascular amyloids (Tokuda et al. 1991). The severity of CTE can be related to the extent of NFTs and their distribution, as found post-mortem (McKee et al. 2013). At its early stages, CTE neurodegeneration is diagnosed through the perivascular accumulation of the NFTs below the sulci depth, which is usually observed in the frontal lobe (McKee et al. 2016). Greater severities are realized when NFTs are widespread and seen in high densities in cortical areas, thalamus, hypothalamus, and other brain regions (McKee et al. 2013; McKee et al. 2016). Considering the correlation of CTE localization with the negative curvature sulci during the early stages of neurodegeneration (McKee et al. 2016), the importance of understanding the effects of complex brain geometries and heterogeneities becomes apparent.

The human brain is both heterogeneous (white/gray matter, fractal vasculature) and anisotropic (*e.g*., radial cortical organization, oriented fiber tracts). Gray matter is located on the outer surface of the brain and contains numerous cell bodies and neuronal somas. The gray matter surrounds the white matter that mostly consists of myelinated axon tracts. In addition, to accommodate the large cortical sheet in a limited volume, the neocortical gray matter is more folded in large-brained animals (Allman). The gyri and sulci are the convex and concave folds of the cerebral cortex, respectively. The gyri often abut the inside surface of the cranium, except in major fissures or involutions such as the insula. The major cortical features (*e.g*., the Sylvian fissure and central sulcus) are readily identifiable with some detailed variation in the folding from person to person. The extent of brain convolutions also vary due to malformations, such as lissencephaly and polymicrogyria that enhance or reduce folding on the brain surface (Allman). Furthermore, the brain is encased in a multi-layered, fibrous structure that includes the pia, arachnoid, and dura mater, of which the pia mater is on the surface of the brain and follows its folds. An understanding of these features is then required to decide what features to include in a head model that can be achieved using computational approaches.

Computational head models (Horgan & Gilchrist 2003; Takhounts et al. 2008; Ho et al. 2009; McAllister et al. 2012; Yang et al. 2014; Ghajari et al. 2017; Ganpule et al. 2017; Fernandes et al. 2018) informed from material constitutive models (Mendis et al. 1995; Donnelly et al. 1997; Arbogast & Margulies 1999; el Sayed et al. 2008; Hosseini Farid et al. 2017; Hosseini-Farid et al. 2019) with various injury criteria (Newman 1986; Newman & Shewchenko 2000; Kimpara & Iwamoto 2012; Takhounts et al. 2013) are used to predict injury under various boundary conditions with some frameworks developed to specifically target injury (Horstemeyer et al. 2019). Some of these injury criteria implemented on macroscale head models included significant uncertainty when compared to neurological injury in real-world injury scenarios (Marjoux et al. 2008). Computational head models generally lack sulci in their brain geometries, which has been shown experimentally to produce heterogeneous von Mises strain fields with greater magnitudes close to sulci (Lauret et al. 2009). Macroscale head models that include sulci also have difficulty capturing the localization of such stress and strain concentrations (Ho & Kleiven 2009; Song et al. 2015). More recently, researchers have used mesoscale brain tissue FE models with nonlocal mechanical damage to reproduce the localization of tau agglomarates under the sulcus base during shear shearing of the brain tissue (Noël & Kuhl 2019). More detail on FE models looking at CTE and their comparison with the current work is presented in Appendix E.

The contribution herein is a multiscale FE approach (motivated from a multiscale paradigm shown in Figure 1 (Murphy et al. 2016; A.H. Bakhtiarydavijani et al. 2019; Murphy et al. 2019; A. Bakhtiarydavijani et al. 2019). The multiscale paradigm depicted in Figure 1 tracks biomechancial features of brain injuries through different length scales that identify the most important variables at each length scale and bridges the information between scales. In the current work, the effects of local geometrical variations arising from sulci that are difficult to capture in macroscale FE head impact models are studied using mesoscale models considering the initial stress wave of impact. For this purpose, a set of mesoscale 2D brain FE simulations for which the boundary conditions were garnered from a previously validated macroscale FE whole head simulation, as shown in Figure 2, (Johnson et al. 2016) were used to evaluate the importance of the structural complexities at the mesoscale. The macroscale FE simulations represented the frontal head impact of an American football player wearing a helmet that was statistically the most significant head impact scenario in American football (Crisco et al. 2010). The mesoscale 2D FE simulations represent the frontal, occipital, temporal, and parietal lobes. These simulations consider structural and geometrical complexities, including gray matter, white matter, pia mater, and sulci, while also considering the brain-CerebroSpinal Fluid (CSF) interface properties. The boundary conditions of these mesoscale simulations are informed from the macroscale FE simulations. The resulting stress waves in the mesoscale 2D FE simulations are then discussed and compared to clinical studies of CTE, thus showing how this multiscale modelling approach can provide more accurate results without significantly increasing computational costs.

**Figure 1.**
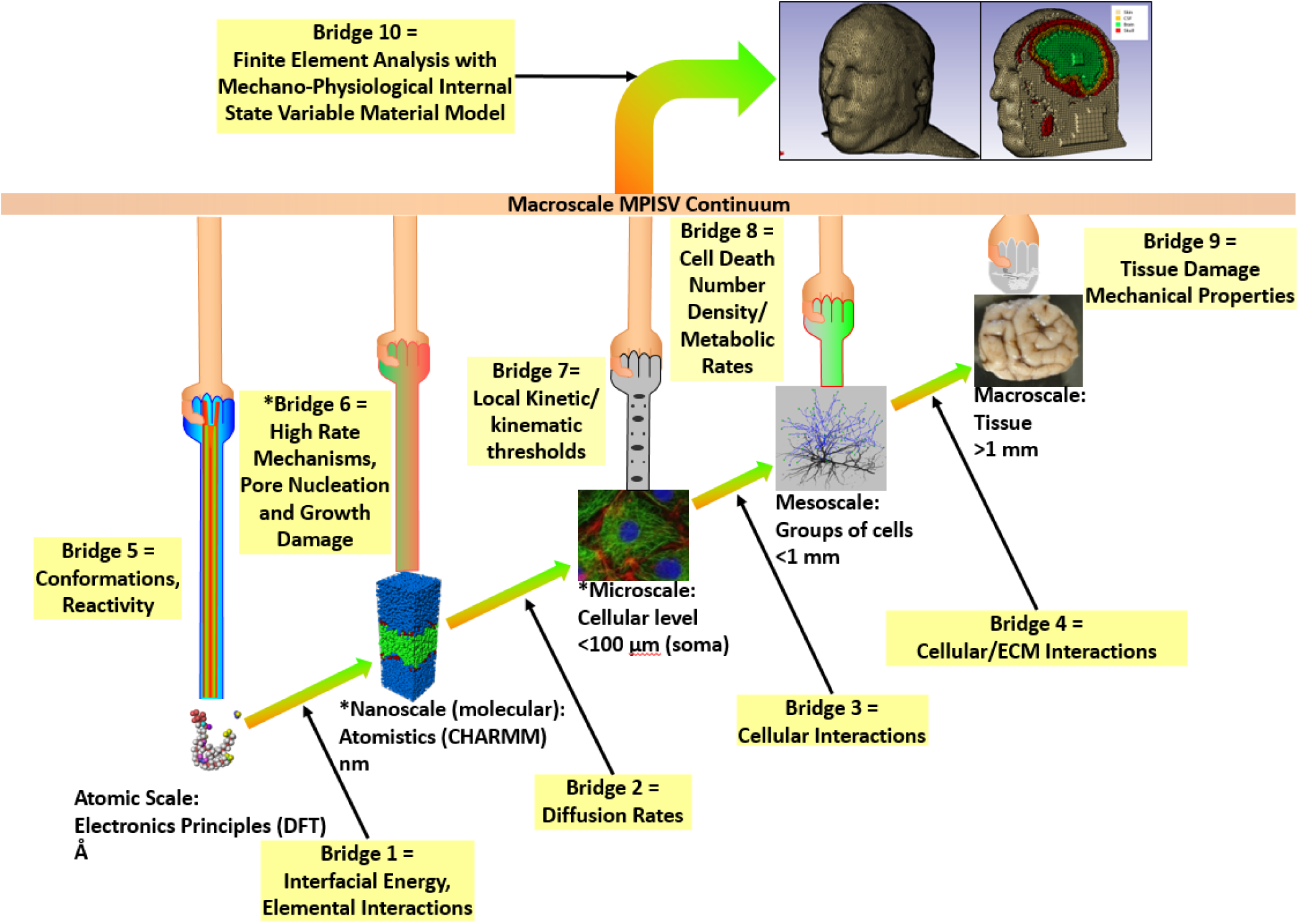
Multiscale approach to modeling neuronal mechanical behavior as it relates to the mechano-physiological internal state variables (MPISV). Modified from Murphy *et al.* (2016).

**Figure 2.**
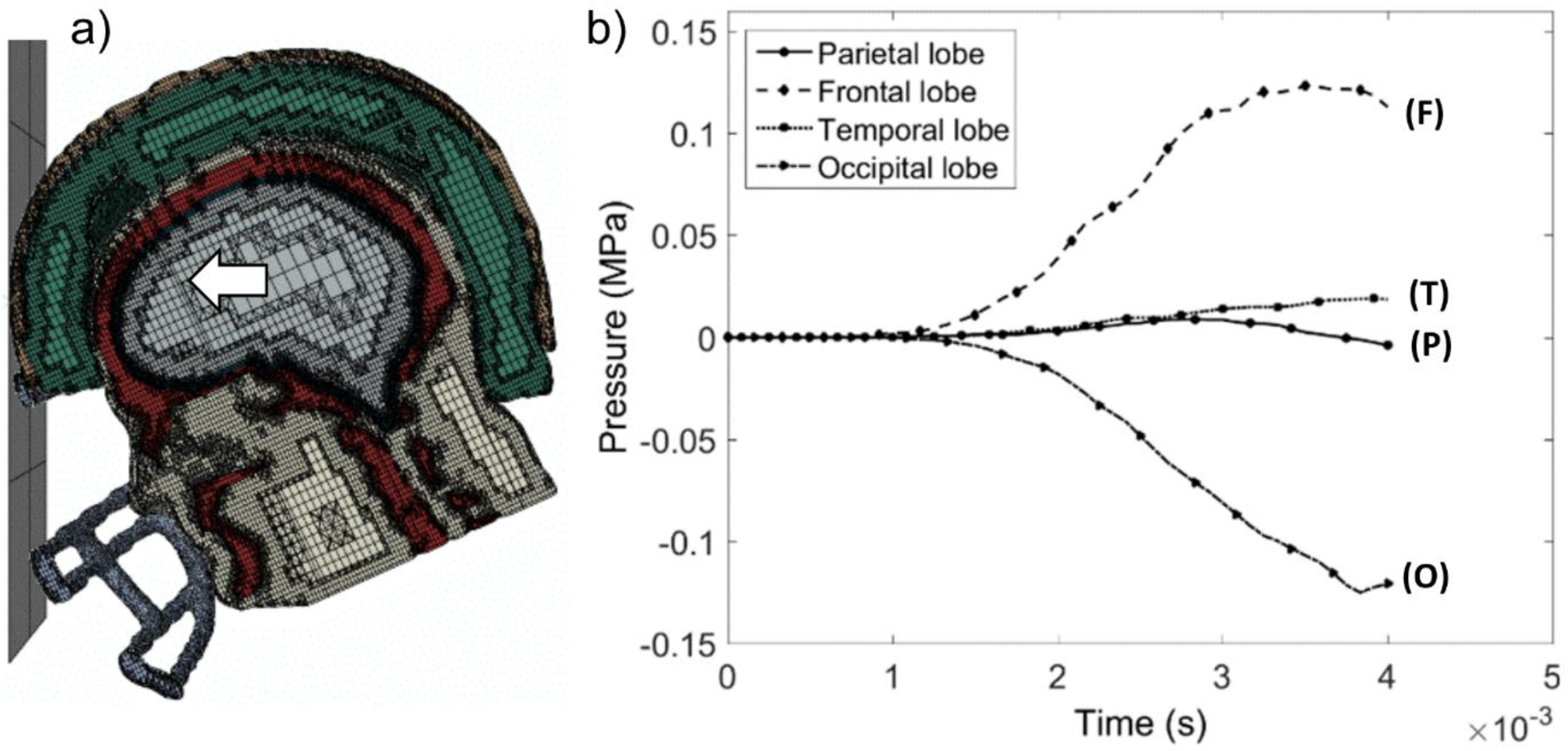
a) Sagittal-cut plane view of the macroscale human head blunt impact finite element model showing the frontal head impact of a helmeted American football player with the white arrow showing the head movement direction and b) the resulting pressure-time histories extracted from the surface of the brain lobes.

## Methods

The macroscale whole head FE simulation included an American football player wearing a helmet. This simulation included the facemask, helmet shell, helmet liner, flesh, cortical and cancellous bone, cerebrospinal fluid (CSF), and the brain. The model was meshed with nearly million elements (Johnson et al. 2016) and validated to Nahum et al. (Nahum et al. 1977). All simulations were performed in the explicit FE software Abaqus/ExplicitTM (Dassault Systemes, Rhode Island, USA). The macroscale model with the Frankfort plane rotated forward, underwent a mid-sagittal head impact (Figure 2a) at 6 m/s. Average pressure-time histories, from four to five nodes that had insignificant (< %5) variation for peak pressure, were extracted from the surface of the occipital lobe, parietal lobe, temporal lobe, and frontal lobe of the frontal head impact simulations as presented in Figure 2b. These pressure-time histories were then applied uniformly to their respective mesoscale 2D FE model described in the next section.

Two sets of mesoscale 2D FE simulations were used to evaluate (1) the effects of various material heterogeneities (Figure 3) and (2) the effects of sulci and gray matter in different brain lobes (Figure 4), when considering the pressures experienced during the frontal head impact. Table 1 presents a list of the FE simulations with the included anatomical features and brain lobes. The base simulation (Figure 3a) included two 25 mm deep, 1 mm wide sulci that were 16 mm apart. The brain tissue was differentiated into gray and white matter, where the gray matter had a 3 mm thickness. Figure 3b shows the same sulci geometry but did not differentiate gray matter from white matter. Figure 3c shows differentiated gray-white matter but lacked sulci. Figure 3d was made without sulci and did not differentiate between the gray and white matter. The effects of the brain-CSF interface were studied by varying the tangential friction coefficient as the following: 0 (frictionless), 0.075, 0.15, 0.30, and 1 (tied) in Figure 3a. A simulation with pia mater was also employed with a 0.1 mm one element thick viscoelastic membrane implemented, as shown in Figure 3a between the CSF and the brain. Also, different lengths of a sulcus were analyzed as well. After identifying the most significant heterogeneities in the brain, the 2D geometry presented in Figure 3a was rotated to create geometries representing different brain lobes, as shown in Figure 4a. Here, the infinite boundary element was placed opposite to the incoming pressure wave (Figure 4b through Figure 4e.). All simulations were performed in the explicit FE software Abaqus/ExplicitTM. These simulations modelled the first 4 ms of impact using a dynamic explicit step with no mass scaling. Forces were introduced as a uniform pressure on the loading surface.

**Table 1.**
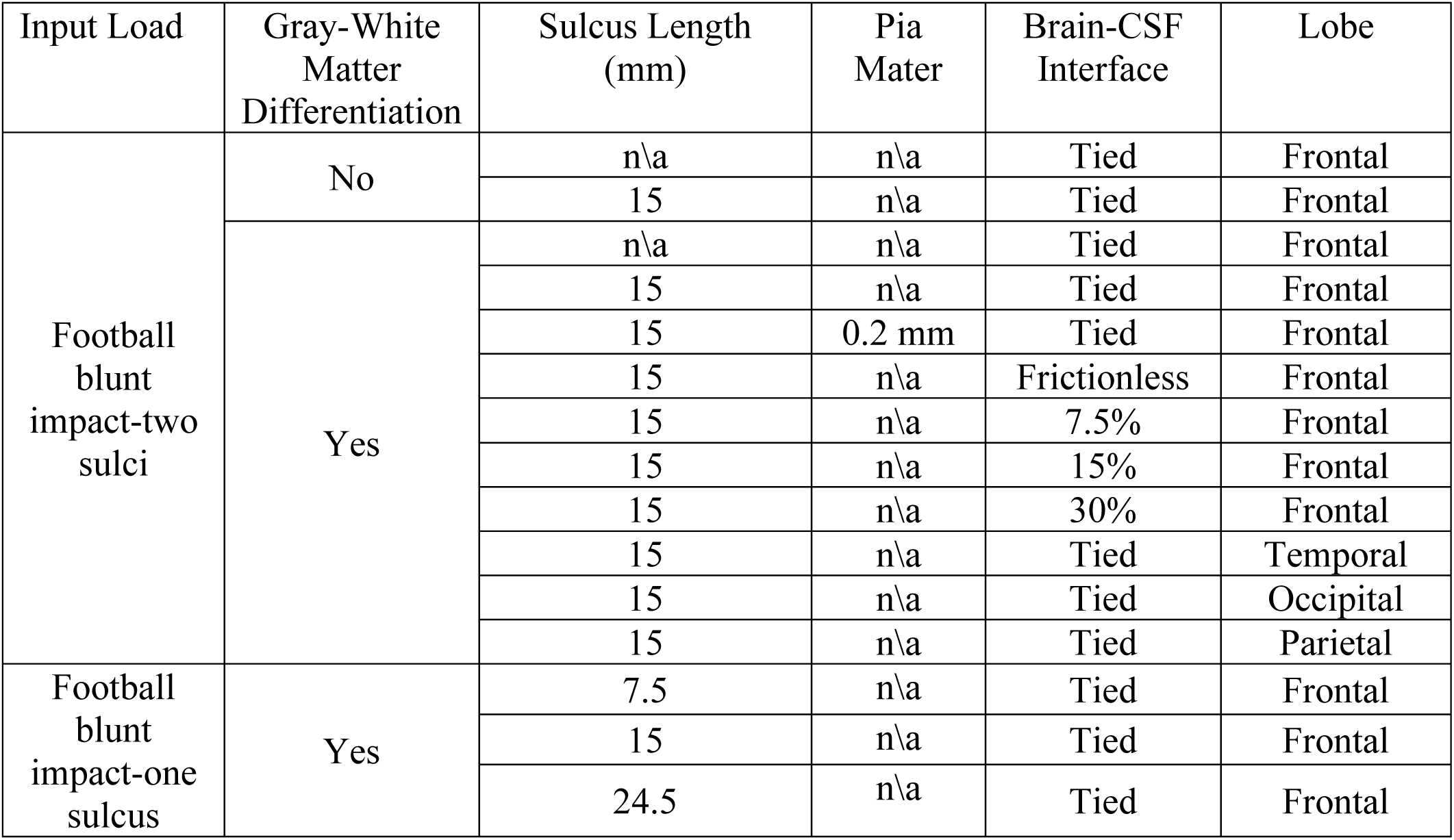
List of the two-dimensional (2D) finite element (FE) models considering the anatomical features, locations, and impact scenario.

**Figure 3.**
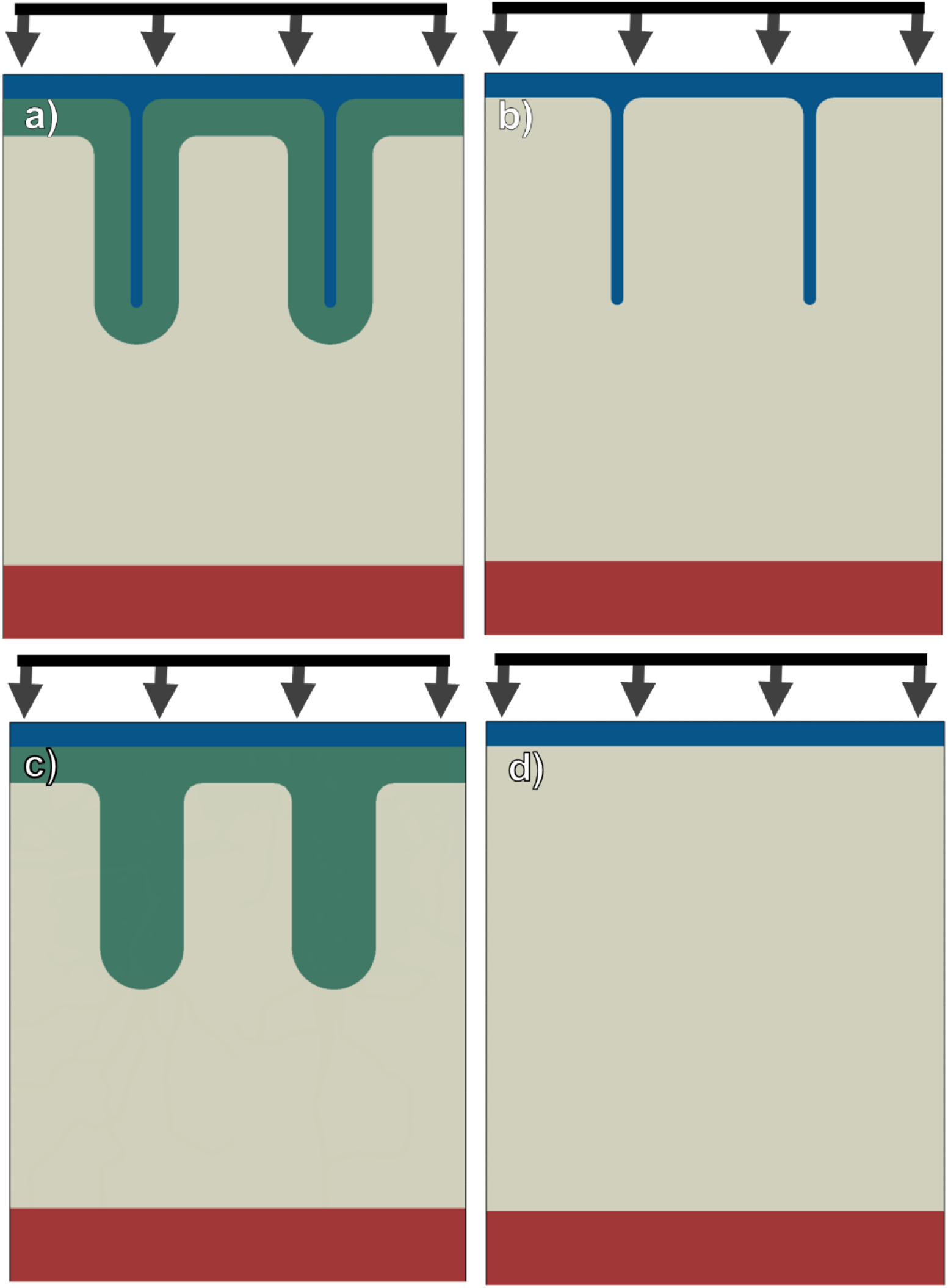
Mesoscale 2D Finite Element (FE) Geometries used to study the effects of structural complexities under the frontal American football player blunt head impact where blue, green, gray, and maroon represent the CSF, gray matter, white matter, and infinite boundary element, respectively. These FE geometries include: a) gray matter, white matter and sulci, b) sulci and homogeneous brain, c) gray matter and white matter, and d) homogeneous brain.

**Figure 4.**
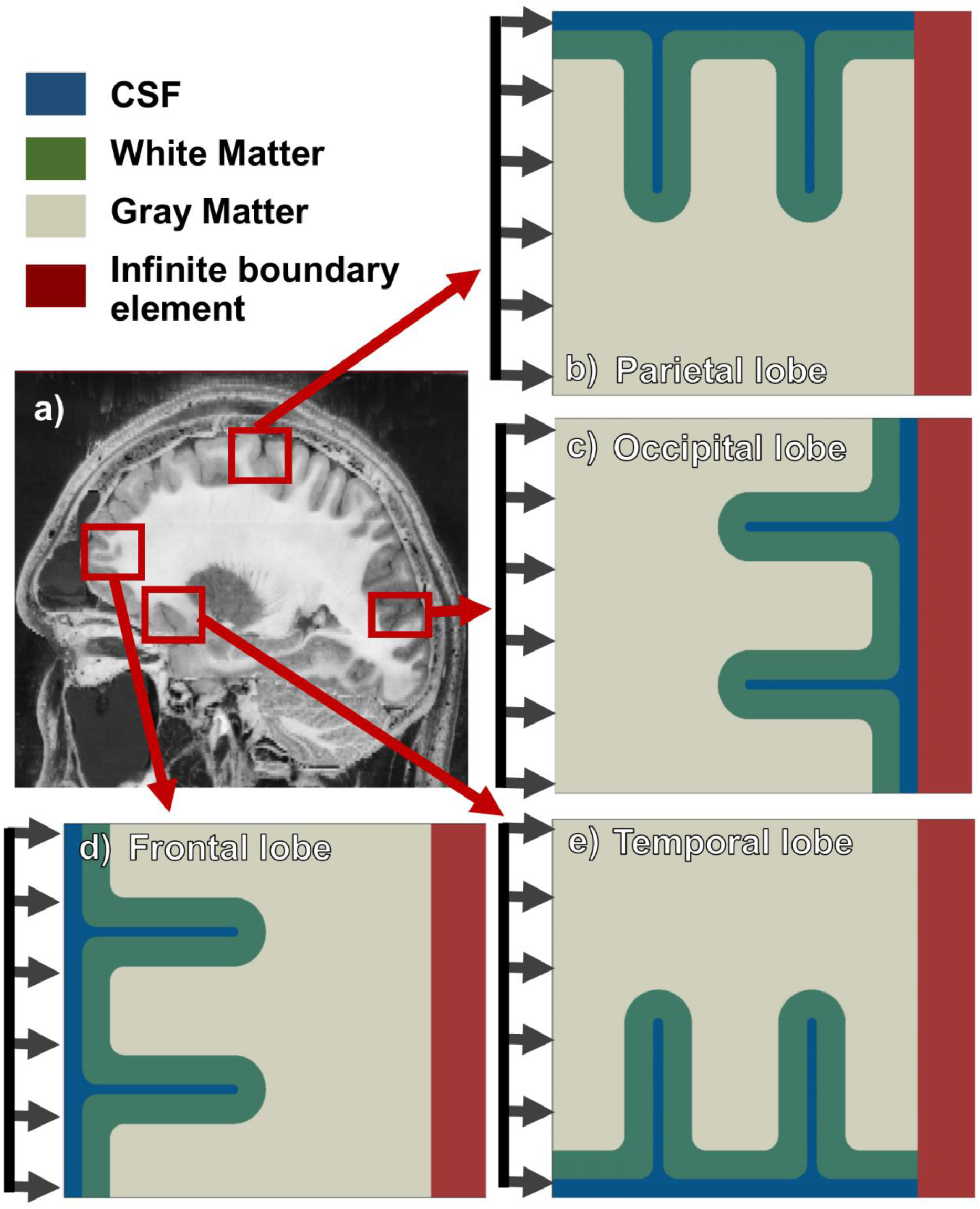
a) Magnetic Resonance Image (MRI) shown as a representative for the parasagittal plane of the human brain with locations highlighted correlating to two-dimensional finite element models of b) the Parietal lobe, c) Occipital lobe, d) the frontal lobe, and e) the temporal lobe with uniform loading direction for the frontal head impact scenario.

### Material Properties

The gray and white matter material properties were defined using the VUMAT MSU TP 1.1 material model that is an elasto-viscoplastic Internal State Variable (ISV) material model initially developed by (Francis et al. 2014) and used to model brain tissue (Prabhu et al. 2011; Johnson et al. 2016) (presented in Appendix A along with the calibrated constants from Johnson *et al.* (Johnson et al. 2016) presented in Table A.1). The CSF was treated as an incompressible elastic material and the pia mater as a viscoelastic material (Aimedieu & Grebe 2004). The constants for these materials are presented in Table 2.

**Table 2.**
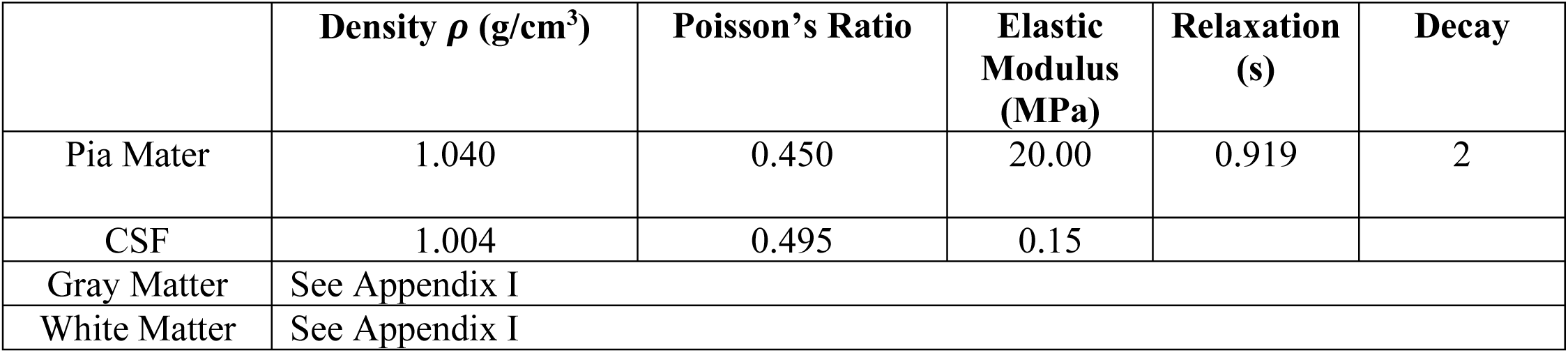
Material properties of the head for the mesoscale finite element simulations.

|  | **Density $\rho$ (g/cm<sup>3</sup>)** | **Poisson's Ratio** | **Elastic Modulus (MPa)** | **Relaxation (s)** | **Decay** |
| --- | --- | --- | --- | --- | --- |
| Pia Mater | 1.040 | 0.450 | 20.00 | 0.919 | 2 |
| CSF | 1.004 | 0.495 | 0.15 |  |  |
| Gray Matter | See Appendix I |  |  |  |  |
| White Matter | See Appendix I |  |  |  |  |

### Boundary Conditions and Mesh

All mesoscale 2D FE simulations used symmetric boundary conditions for boundaries adjacent to the loading surface, and an infinite boundary opposite of the loading surface that prevented pressure wave reflections at that surface. This boundary condition setup showed a good correlation with an extended geometry model that produced zero stress reflections. This was compared to simulations with three symmetric boundaries and three infinite boundary conditions, which gave unrealistic stresses and strains (Table 3). Hence, the single infinite boundary setup was used in all simulations. All simulations used reduced quadratic plane strain elements (Abaqus CPE4R) for the brain and infinite boundary elements (Abaqus CINPE4) for the infinite boundary. A mesh convergence study was performed (Figure 3a.) with 280,000; 30,000; and 9000 elements (or an element size of less than 0.5 mm for the latter mesh). The model with 9000 elements generated a result that was within 5% standard error of the 280,000-element simulation when considering the average and peak pressure and, as such, was used for all simulations. To achieve more detailed field output gradients at the depth of the sulcus, the elements decreased to 0.1 mm below the sulcus.

**Table 3.** Validating boundary condition elements in mesoscale 2D Finite Element (FE) simulations of the frontal lobe during the frontal head impact of a football player. The table presents the localized stresses and strains below the sulcus and the average pressure of the whole model.

| **Model** | **Average element length (mm)** | **$\sigma_{vm}$ (kPa)** | **Peak Pressure (kPa)** | **$\epsilon_1$** | **Average pressure (kPa)** |
| --- | --- | --- | --- | --- | --- |
| Periodic/ infinite boundary element model | 0.5 | 17.93 | 130.0 | 0.02355 | 115.25 |
| Extended geometry model | 0.5 | 16.70 | 136.5 | 0.02556 | 119.74 |
| All periodic boundary model | 0.5 | 285 | 106.8 | 0.1802 | 76.25 |

## Results

Figure 5 presents the von Mises stress and pressure in the frontal lobe during the frontal head impact of an American football player at the time of peak loading. The white arrows in Figure 5a show the direction of the pressure waves, while the gray double-headed arrows show the direction of the largest principal stress. Figures 5b and 5c show Expanded views of the localization of pressure and von Mises stress around a sulcus, respectively.

**Figure 5.**
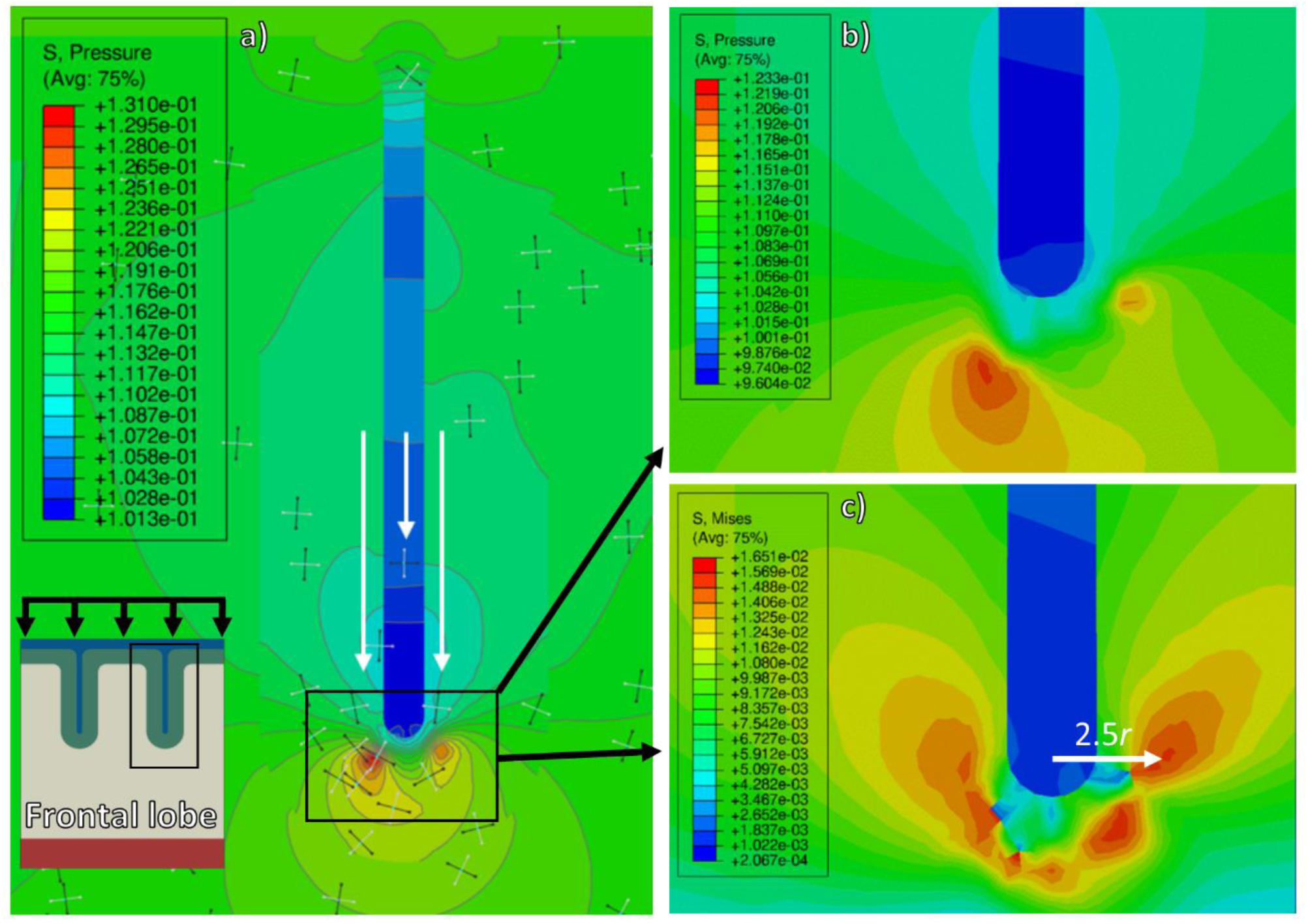
Mesoscale 2D Finite Element (FE) simulations of the Pressure and von Mises stress (MPa) localization below the sulcus in the frontal lobe of a football player experiencing a frontal head impact at 3.85 ms. a) Pressure wave propagation during peak pressures, showing slower wave speeds in the CSF compared to brain. The small gray double headed arrows show the direction of the maximum peak principal stress, and the white arrows show the direction of the stress wave propagation. b) The localization of pressure below the sulcus arises from the geometrical complexity (sulcus), and the interaction of the two pressure waves on either side of the sulcus. c) The von Mises stress localization arising from the pressure field below the sulcus.

Figures 6 and 7 present the von Mises stress and pressure evolution in the frontal lobe, respectively. Here, the four different structural heterogeneities shown in Figure 3 are considered. Figures 6a-c and 7a-c show these temporal evolutions for a simulation that includes sulci and gray-white matter differentiation with 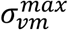 and maximum pressure values of 18.0 kPa and 131.0 kPa, respectively. Figures 6d-f and 7d-f show these values for a simulation that includes sulci and homogeneous brain material with 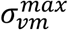 and maximum pressure values of 17.7 kPa and 127.0 kPa, respectively. Figure 6g-i and Figure 7g-i only differentiate gray-white matter but do not include sulci and result in 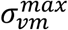 and maximum pressure values of 0.6 kPa and 122.7 kPa, respectively. Figure 6j-l and Figure 7j-l consider a homogenous brain material without sulci and result in 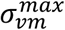 and maximum pressure values of 0.5 kPa and 122.5 kPa, respectively.

**Figure 6.**
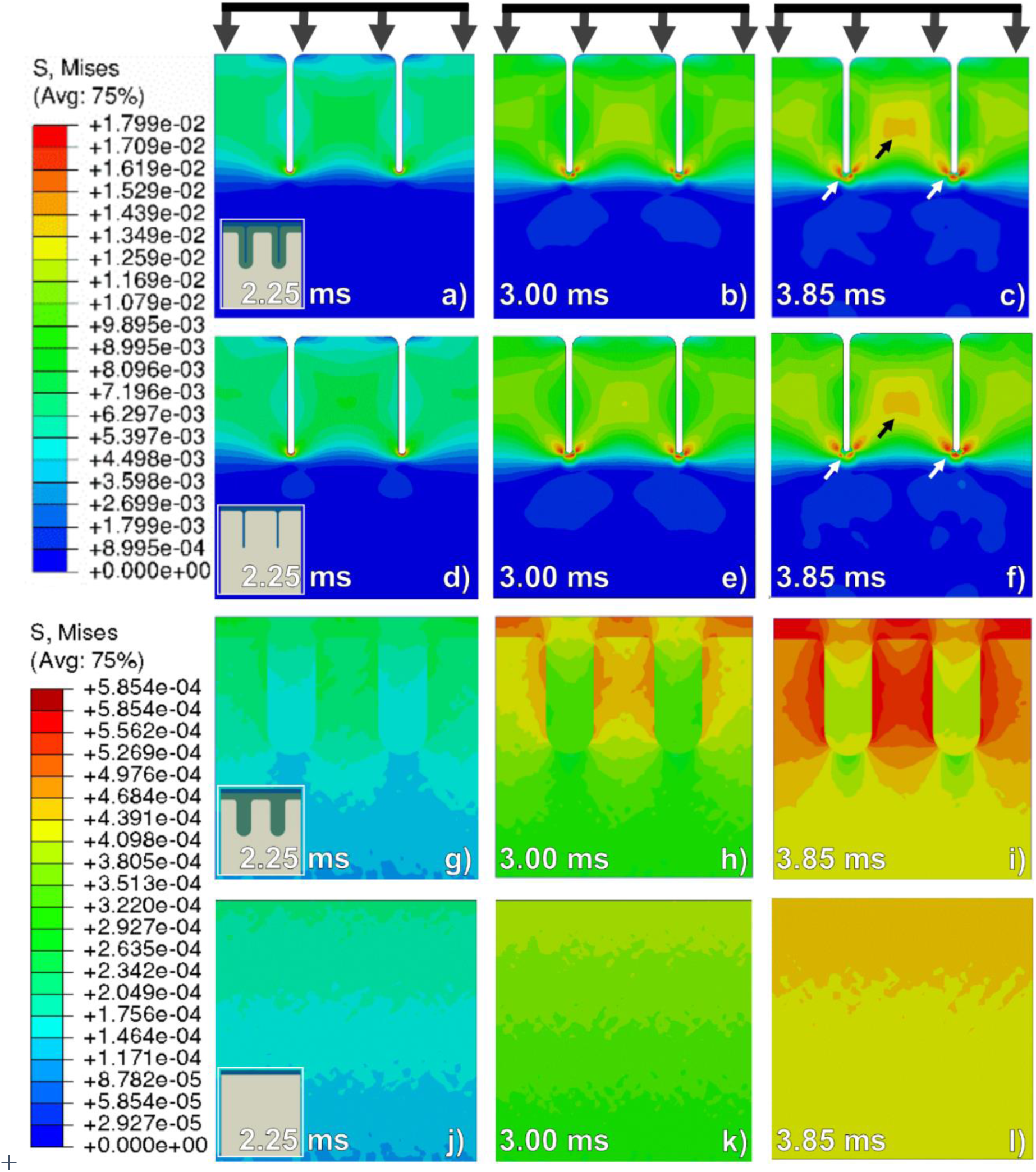
Von Mises stress (MPa) evolution in simulations with different geometry and structural compositions. All simulations were loaded from the top with the pressure-time histories observed in the frontal lobe during the football player head impact (Fig. 3), where the three columns, from left to right, present resulting von Mises stress contours at 2.25, 3.00, and 3.85 ms, respectively. These models include (a) – (c) sulci and differentiated gray-white matter; (d) – (f) sulci and homogeneous brain; (g) – (i) differentiated gray-white matter without sulci; and (j) – (l) homogenous brain with no sulci. White and black arrows point to the location of the von Mises stress localizations arising from the near and far-field effects of sulci, respectively. The CSF and the infinite boundary are not shown for visual clarity.

**Figure 7.**
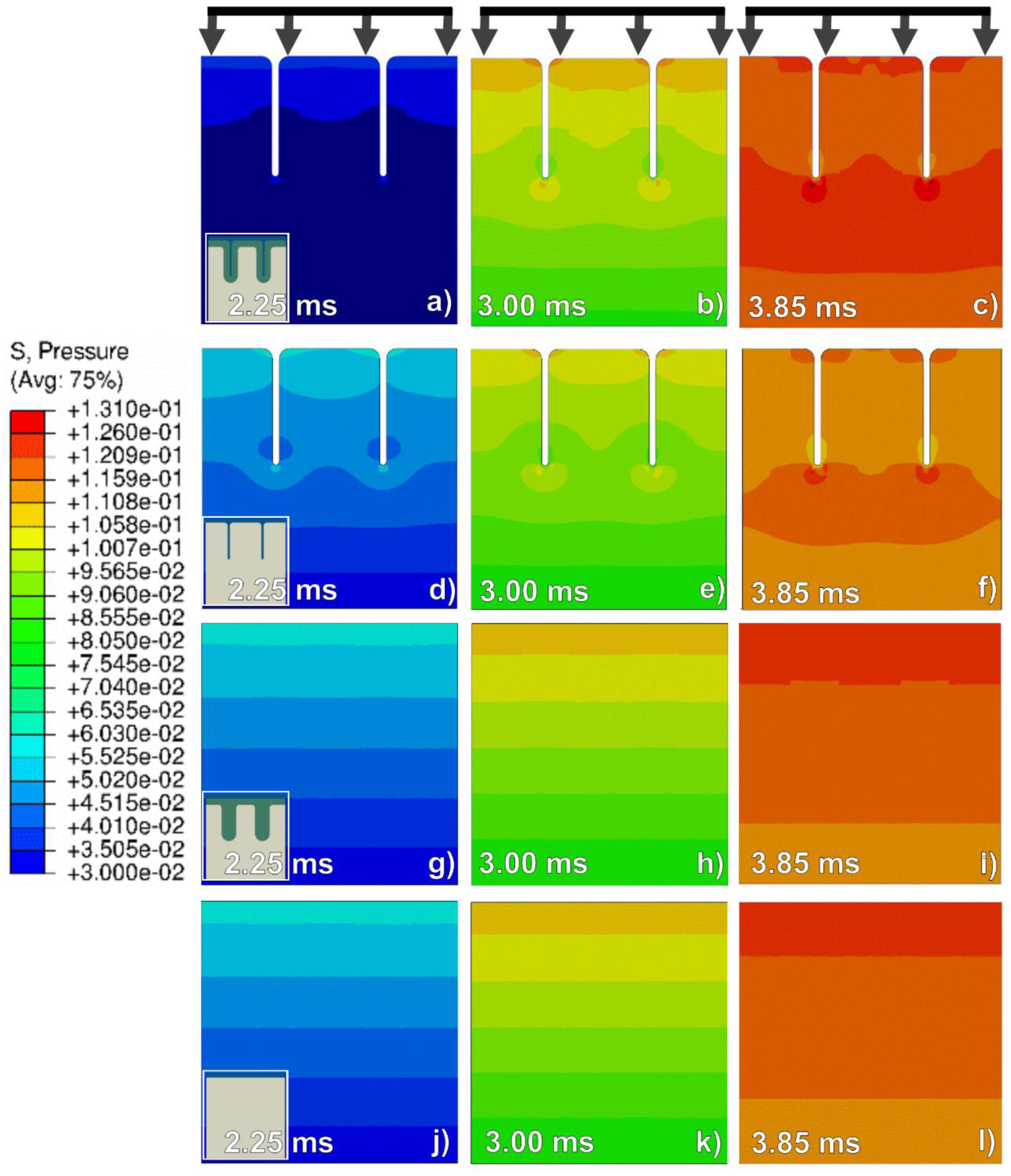
Pressure (MPa) evolution due to different structural complexities in mesoscale 2D finite element simulations. All simulations were loaded from the top with the pressure-time histories observed in the frontal lobe during the football player head impact (Fig. 3), where the three columns, from left to right, present resulting pressure contours at 2.25, 3.00, and 3.85 ms, respectively. These models include (a) – (c) sulci and differentiated gray-white matter; (d) – (f) sulci and homogeneous brain; (g) – (i) differentiated gray-white matter with no sulci; and (j) – (l) homogenous brain with no sulci. The CSF and the infinite boundary are not shown for visual clarity.

Figure 8 shows the von Mises stress (Figure 8a-c) and pressure (Figure 8d-f) that developed under a 15 mm sulcus with the maximum values of 17.8 kPa and 130.9 kPa, respectively. The effects of the sulcus length on the 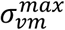 during peak applied pressure are shown in Figure 8g-i, with the 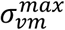 increasing from 4.2 kPa to 6.8 kPa with increasing sulcus length.

**Figure 8.**
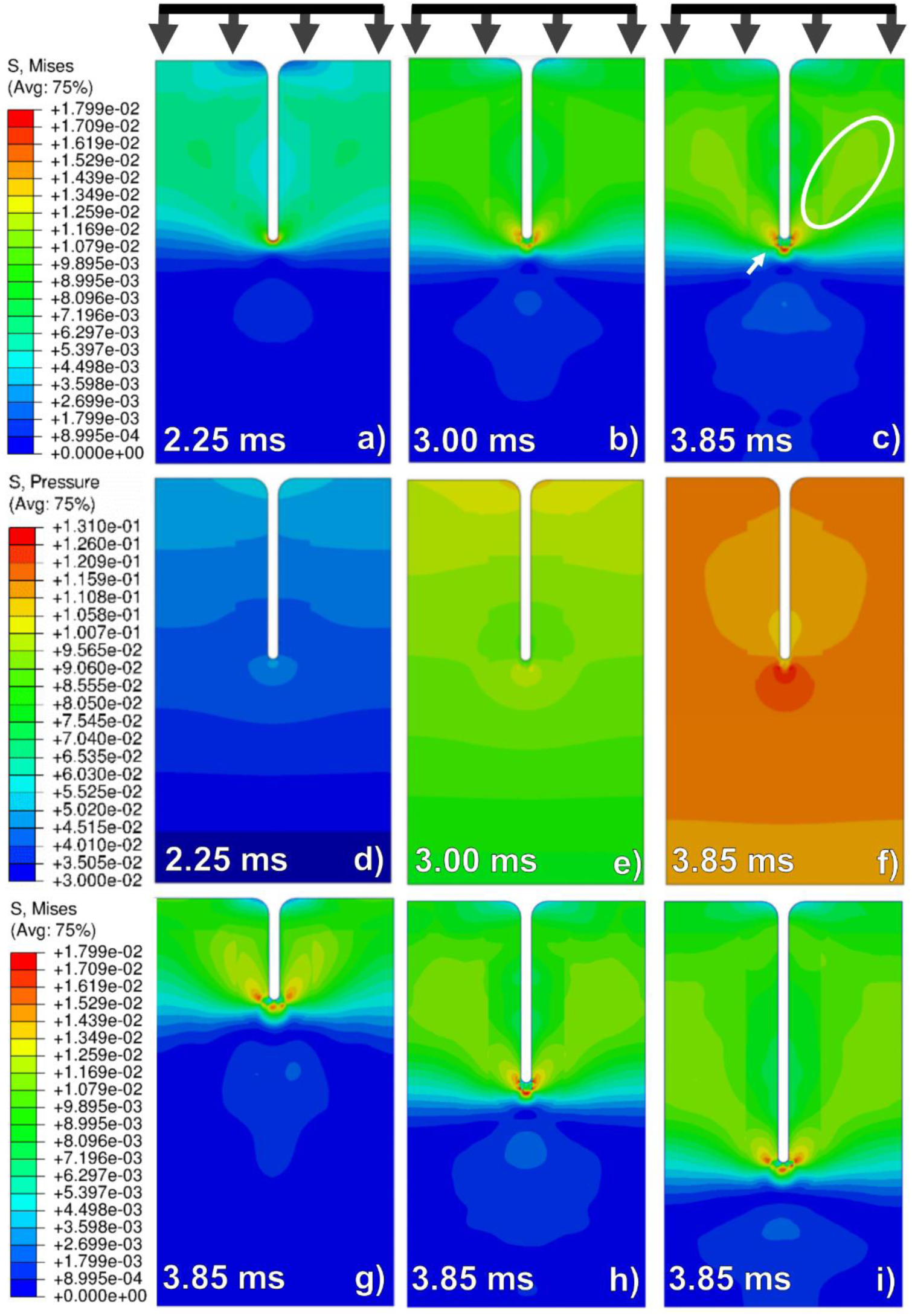
Mesoscale 2D finite element simulations of the frontal lobe with 14.5 mm sulcus during the frontal head impact of a football player simulation illustrating the time evolution of the von Mises stress (MPa) and pressure (MPa) contours at (a) and (d) 2.25 ms, (b) and (e) 3.00 ms, and (c) and (f) 3.85 ms. The von Mises stress contours are then presented at peak applied pressure for different sulcus lengths: (g) 7.5 mm, (h) 14.5 mm, and (i) 24.5 mm. The white arrow shows the von Mises stress localization arising from the inclusion of a sulcus with the von Mises stress band circled in white. The CSF and the infinite boundary are not shown for visual clarity.

Figures 9 and 10 show the von Mises stress and pressure histories in the four different brain lobes (Figure 4b-e). Figure 2b shows the applied pressure boundary condition for each mesoscale simulation. The four rows in these figures are the evolution contours in the frontal, parietal, temporal, and occipital lobes, respectively. When sulci are parallel to the incoming pressure wave, both the von Mises stress and pressure initially localize below the sulci and move inward into the sulci depth as time progresses (Figures 9c and 9l). When the sulci are perpendicular to the incoming pressure wave, stress concentrations localized below the sulci (Figures 9f and 9j). The pressure profiles also show peak stress concentrations below the sulci for all simulations. Table 4 summarizes the average and maximum von Mises stresses and pressures.

**Table 4.**
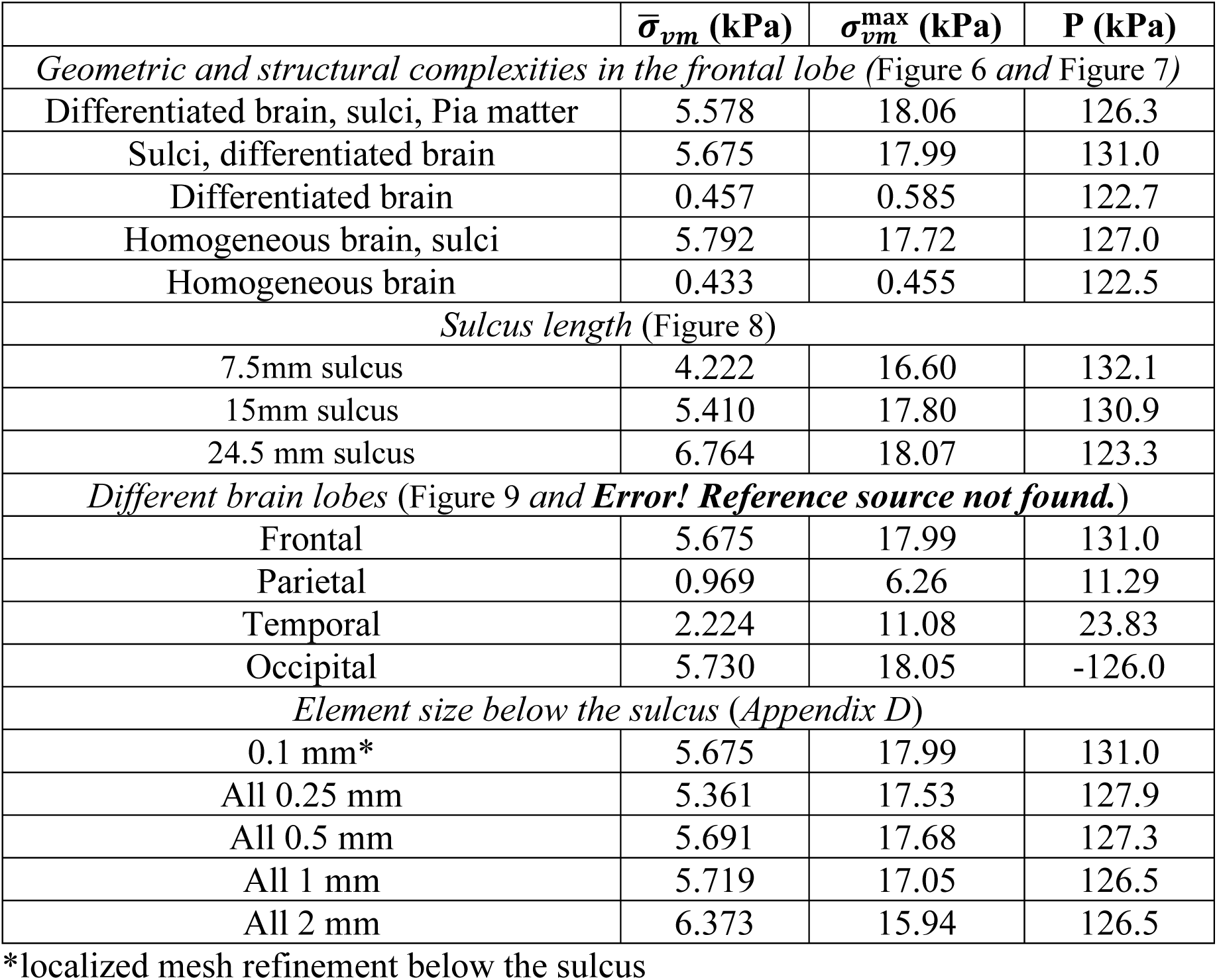
Summarized average von Mises stress 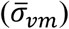, maximum von Mises stress 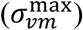, maximum pressure (P) observed in the mesoscale finite element simulations.

| | $\bar{\sigma}_{vm}$ (kPa) | $\sigma_{vm}^{\max}$ (kPa) | P (kPa) |
| --- | --- | --- | --- |
| *Geometric and structural complexities in the frontal lobe (Figure 6 and Figure 7)* |  |  |  |
| Differentiated brain, sulci, Pia matter | 5.578 | 18.06 | 126.3 |
| Sulci, differentiated brain | 5.675 | 17.99 | 131.0 |
| Differentiated brain | 0.457 | 0.585 | 122.7 |
| Homogeneous brain, sulci | 5.792 | 17.72 | 127.0 |
| Homogeneous brain | 0.433 | 0.455 | 122.5 |
| *Sulcus length (Figure 8)* |  |  |  |
| 7.5mm sulcus | 4.222 | 16.60 | 132.1 |
| 15mm sulcus | 5.410 | 17.80 | 130.9 |
| 24.5 mm sulcus | 6.764 | 18.07 | 123.3 |
| *Different brain lobes (Figure 9 and **Error! Reference source not found.**)* |  |  |  |
| Frontal | 5.675 | 17.99 | 131.0 |
| Parietal | 0.969 | 6.26 | 11.29 |
| Temporal | 2.224 | 11.08 | 23.83 |
| Occipital | 5.730 | 18.05 | -126.0 |
| *Element size below the sulcus (Appendix D)* |  |  |  |
| 0.1 mm* | 5.675 | 17.99 | 131.0 |
| All 0.25 mm | 5.361 | 17.53 | 127.9 |
| All 0.5 mm | 5.691 | 17.68 | 127.3 |
| All 1 mm | 5.719 | 17.05 | 126.5 |
| All 2 mm | 6.373 | 15.94 | 126.5 |

\*localized mesh refinement below the sulcus

**Figure 9.**
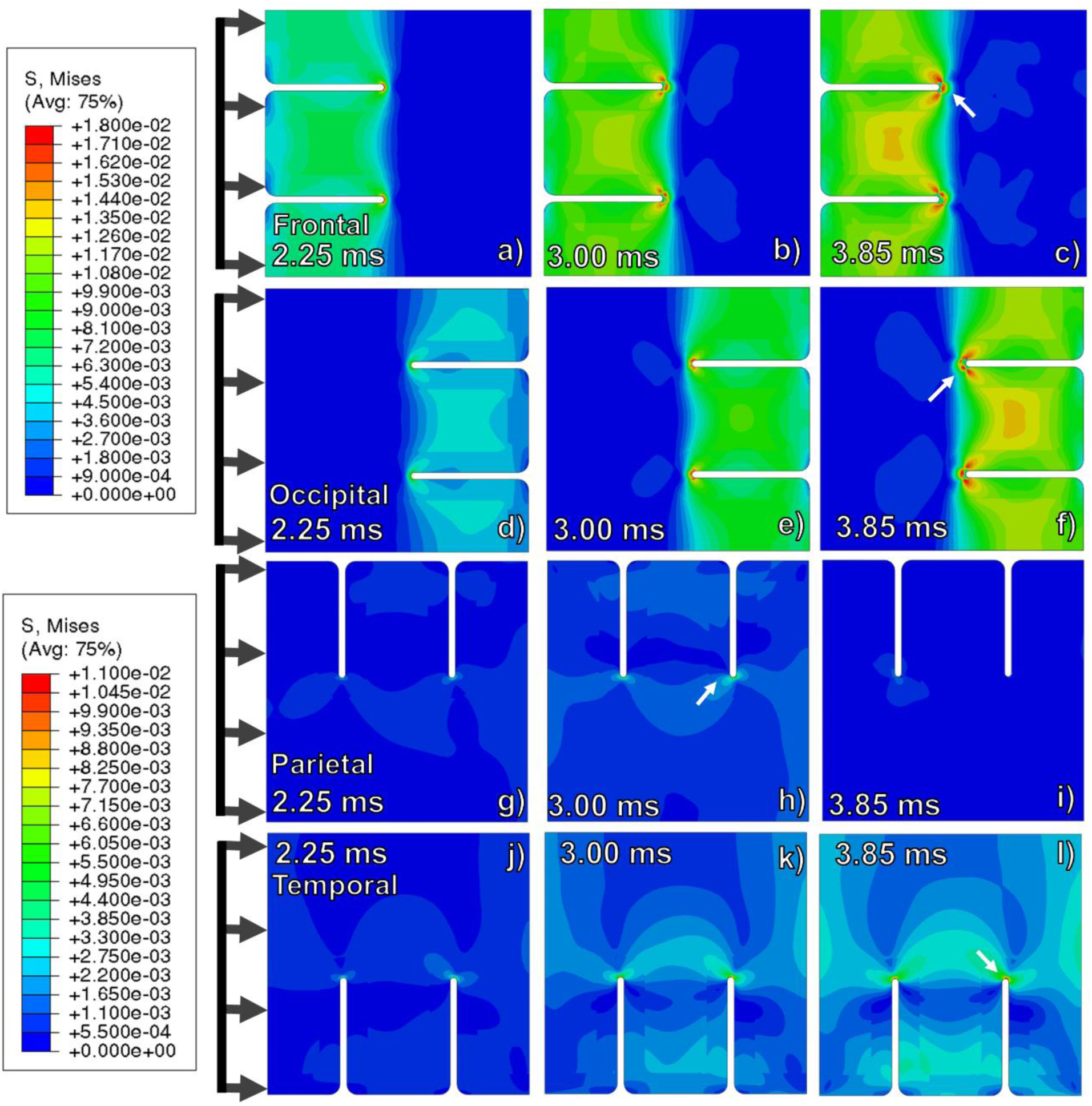
Mesoscale 2D finite element simulations of the four different brain lobes illustrating the von Mises stress (MPa) evolution experienced during the frontal head impact of a helmeted football player. The three columns from left to right represent von Mises stress contours at 2.25, 3.00, and 3.85 ms, respectively. These simulations represent (a) – (c) the frontal lobe, (d) –(f) the occipital lobe, (g) – (i) the parietal lobe, and (j) – (l) the temporal lobe, with the white arrows showing the location of peak von Mises stress. The load is applied from the left side. The CSF and the infinite boundary are not shown for visual clarity.

## Discussion

The current work highlights the effectiveness of mesoscale FE simulations in capturing the relevant length scale geometrical complexities that are generally unattainable with macroscale FE simulations. The present 2D mesoscale FE simulations garner their boundary conditions from 3D macroscale FE simulations and are used to study the brain’s geometric complexities and its correlation with CTE biomechanics in American football player head impact scenarios. Based on the aforementioned results, we observed that CTE initiation time and location are determined by two things: 1) structural heterogeneities, like sulci and gyri, that give rise to stress concentrators and gradients in the frontal, occipital, parietal, and temporal brain lobes and 2) the difference between the mechanical properties of the CSF and the brain. Thus, the brain’s geometries and material heterogeneities were examined through mesoscale 2D FE simulations to determine their effects on the stress state and the limits of which can indicate injury (Chatelin et al. 2011). Specifically, the mesoscale 2D FE simulations accommodate the anatomical geometries of the brain’s sulci and gyri, gray-white matter interface, and pia mater with a reasonable computational cost. This greater resolution provides a more detailed description of the stress wave propagation and the associated stress concentrations around the sulci. The locations of the stress concentrations correlated well with the sites of tau protein accumulation in Stage I CTE (McKee et al. 2016). Although the correlation between CTE and head impact has been clearly shown (Mez et al. 2017), the mechanism of CTE is still under study (Katsumoto et al. 2019). These stress concentrations were not captured in previous macroscale simulations (Ho & Kleiven 2009; Song et al. 2015; Johnson et al. 2016). But, mesoscale brain models of brain tissue powered by elastic modulus softening, showed the strain energy localization under large shear strains perpendicular to the sulci (Noël & Kuhl 2019).

When studying the von Mises stress (*σ*_*vm*_) evolution for a frontal head impact, the peak *σ*_*vm*_ below the sulci for the frontal and occipital lobes were 18.0 kPa and 19.1 kPa (Table 4), respectively. In these brain lobes, *σ*_*vM*_ initiated at the sulci end at the gray matter-CSF interface (Figures 9a, 9b, 9j, and 9k) and, with time, increased inward into the gray matter at the depth of the sulci forming a “cup shape” von Mises stress concentration region at the sulcus depth (Figures 9c and 9l). At the same time, shear bands initiated from the sulci base that extended into the white matter, overlapping with those of other sulci to produce a secondary area of stress concentration when more than one sulcus was present.

Investigation of stress waves traversing in the vicinity of sulci showed that they affected the internal stresses in two different regimes, described here as near-field and far-field. In the near-field, a sulcus produced local pressure and von Mises stress concentrations (white arrows in Figure 8c, Figures 6c and 6f compared to Figures 6i and 6l) at the sulci depth. This arose due to the impedance mismatch between the CSF and the brain tissue (Figure 5), causing the stress waves to travel slower in the CSF than in the brain tissue. As the faster pressure waves travelled in the brain matter adjacent to the sulcus and wrapped around the sulcus end (Figure 5b), they interacted with each other to form pressure and von Mises stress localizations (Figure 5c). This stress wave interaction was further complicated by the slower stress waves entering the gray matter through the CSF. The mismatch between the wave speeds in the brain and CSF further disrupted the pressure wavefront and intensified the maximum pressure localization near the sulci end (Figure 7c compared to Figure 7l). When sulci were present the 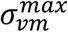 increased by a little more than 39 times from 0.46 kPa to 18.0 kPa (Figure 6l compared to Figure 6f). When comparing the FE simulation results with and without a sulcus, the increase in the maximum pressure was not as significant as that of the von Mises stress (maximum pressure increased by 7% from 122.5 kPa to 131.0 kPa (Figure 7f versus Figure 7l) compared to 39 times). Thus, the simulations support the proposition that shear localization can be the proximal cause of neurodegeneration. Specifically, the near-field shear localization induced by the presence of sulci illustrates (Figure 6) the biomechanics of structural damage that lead to CTE (McKee et al. 2016).

The far-field effects of sulci arise from the inter-sulcus stress field interactions. When a single sulcus was present (Figure 8), shear bands that originated at the sulcus end extended into the brain tissue (Figure 8c, circled white) resulting in 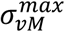 of 12 kPa. When another sulcus, of the same length, was introduced (Figure 6c and 6f, black arrows), the shear bands of the two sulci overlapped to produce a new area of stress localization with a 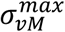 of 14.8 kPa (Figure 9c) that was 23% greater than the 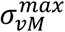 of measured in the one sulcus simulation (12 kPa, Figure 8c, circled white). This far-field stress localization was not observed in scenarios where sulci were perpendicular to the incoming pressure wave (Figure 9h and 9l). The location and existence of this far-field stress localization were found to be dependent on the sulcus length (Figure 8) and orientation (Figure 9) relative to the incoming pressure wave.

A previously published study has shown that large shear strains perpendicular to the sulci can localize the strain energy below the sulcus (Noël & Kuhl 2019). This study had investigated one sulci orientation where the displacament boundary condition were applied perpendicular to the sulcus (Noël & Kuhl 2019). Thus, in order to understand the influence of sulcus orientation to the impending pressure waves, we considered four sulci orientations with respect to the incoming pressure waves due to a frontal impact as observed in the frontal, pariental, temporal, and occipital lobes. This orientation aspect is important to consider, especially during a frontal head impact, which is statistically most significant head impact scenario in American football (Crisco et al. 2010). During a frontal head impact the pressure waves propagate parallel to the sulci in the frontal and occipital lobes, and perpendicular to the sulci in parietal and temporal lobes (McKee et al. 2016) (Figure 4). When studying the 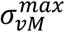 field in these mesoscale models, it was found that the *σ*^max^ in the parallel configuration (frontal and occipital lobes) are significantly greater than that in the perpendicular orientation (temporal and parietal lobes), as presented in Table 4. Considering the locally applied pressures (Figure 2b) in conjunction with the 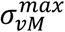 generated in Figure 9 leads to a better understanding of sulci effects on stress localization. During the head impact, sulci parallel to the pressure waves (frontal and occipital lobes) experienced peak pressures that were approximately an order of magnitude greater than the pressures in sulci perpendicular to the stress waves (parietal and temporal lobes). In contrast, the 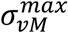 in the perpendicular sulci was only two-thirds of those in the parallel sulci simulations (Table 4). Finally, although not reflected in this work, the frontal head impact may have a smaller effect in the occipital lobe due to wave dispersion or defocusing as it traverses through the brain.

The sulcus length was found to affect the near-field stress concentrations when the impending pressure waves traversed through the CSF and flanked the gray matter of the sulcus. By increasing the sulcus length from 7.5 mm to 24.5 mm, the 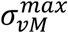 increased below the sulcus end by approximately 9%, from 16.6 kPa to 18.1 kPa (Table 4 and Figure 8g-i). This increase could be related to the increased mismatch between the stress wavefronts traveling in the CSF and gray matter (Figure C1 in Appendix C). In the far-field, the length of the shear bands was proportional to the sulcus length but had only minor effects on the magnitude of the von Mises stress in those bands (Figure 8g-i). The larger area covered by these stress bands means that more brain tissue may be susceptible to injury with longer sulci. These effects may be compounded by brain malformations, such as lissencephaly and polymicrogyria (Budday et al. 2015). Accordingly, the susceptibility of individuals with such conditions to blunt impact head trauma should differ from those without the susceptibility.

## Conclusions

In this work, pressure-time boundary conditions extracted from macroscale human head Finite Element (FE) simulations were used to study stress wave propagations in mesoscale 2D FE simulations of the brain sulcus-gyrus geometries and material heterogeneities in four different brain lobes (frontal, parietal, temporal, and occipital). The following were the salient observations:

- The impedance mismatch of the CSF and the gray matter (Figure 5) produced a variation in the wave propagation speeds in CSF and gray matter that resulted in the complex local interaction of the two stress waves (on both sides of a sulcus) and a wavefront in the CSF at the sulcus end. These stress wave interactions synergistically increased the von Mises (shearing) stress in the near- and far-field of the sulci.
- In the near-field, a sulcus introduces a von Mises stress localization just below the sulcus end (Figure 5.c) that was 39 times greater than those experienced in simulations without sulci (Figure 6.f and 6.i and Table 4). These stress concentrations below the sulci provide a mechanism for the corresponding localized damage (tau protein accumulation) observed during the early stages of CTE in the literature.
- In the far-field, shear stress bands emanate from the sulcal ends (Figure 8.c-white circle) and interact with one another to produce von Mises stress concentrations in the brain tissue (Figure 6.c and f-black arrow) that are 23% greater in magnitude compared to single sulcus simulations. The magnitude and extent of this stress concentration are dependent on the sulci lengths, orientations, and nearest neighbour distances from each other.
- Increasing the sulcus length from 7.5 mm to 24.5 mm increased the resulting von Mises stress concentration by 9% (Figure 8.g-i and Table 4) and produced large von Mises stresses that extended further into the white matter.
- Differentiating between gray and white matter, even with different elasto-viscoplastic constitutive properties, produced small von Mises stress concentrations at the gray-white matter interface when compared to stresses of sulci (Figure 6.c and Table 4). Accounting for sulci presence is essential to modelling the biomechanical causes of CTE.
- Varying the brain-CSF numerical interaction (frictionless, tied, and varying the tangential friction coefficients from 7.5% to 30%) and introducing pia matter into the simulation did not significantly affect the stress wave propagation nor the localization of von Mises stress (Appendix D).

Finally, the current study shows how multiscale FE simulations with whole head FE simulations providing the boundary conditions for lower length scale mesoscale FE simulations can provide a more detailed understanding of regions prone to injury without significant increases in computational costs.

## Acknowledgments

The authors would like acknowledge the Center for Advanced Vehicular Systems (CAVS) at Mississippi State University for supporting this work.

## Ethical approval

Not required

